# A Cyclin A – Myb-MuvB – Aurora B network regulates the choice between mitotic cycles and polyploid endoreplication cycles

**DOI:** 10.1101/449983

**Authors:** Michael D. Rotelli, Robert A. Policastro, Anna M. Bolling, Andrew W. Killion, Abraham J. Weinberg, Michael J. Dixon, Gabriel E. Zentner, Claire E. Walczak, Mary A. Lilly, Brian R. Calvi

## Abstract

Cells switch to polyploid endoreplication cycles during development, wound healing, and cancer. We used integrated approaches in *Drosophila* to determine how mitotic cycles are remodeled into endoreplication cycles, and how similar this remodeling is between developmental and induced endoreplicating cells (devECs and iECs). We found that while only devECs had a dampened E2F1 transcriptome, repression of a Cyclin A - Myb-MuvB - Aurora B mitotic network promoted endoreplication in both devECs and iECs. Cyclin A associated with and activated Myb-MuvB to induce transcription of mitotic genes, with expression of one, Aurora B, being key for mitotic commitment. Knockdown of Cyclin A, Myb, Aurora B, or downstream cytokinetic proteins induced distinct types of endoreplication, suggesting that repression of different mitotic network steps may explain the known diversity of polyploid cycles. These findings reveal how remodeling of a mitotic network promotes polyploid cycles that contribute to development, wound healing, and cancer.

## Introduction

Endoreplication is a common cell cycle variant that entails periodic genome duplication without cell division and results in large polyploid cells (Fox and Duronio, 2013). Two variations on endoreplication are the endocycle, a repeated G/S cycle that completely skips mitosis, and endomitosis, wherein cells enter but do not complete mitosis and / or cytokinesis before duplicating their genome again (Calvi, 2013). In a wide array of organisms, specific cell types switch from mitotic cycles to endoreplication cycles as part of normal tissue growth during development (Orr-Weaver, 2015). Cells also can switch to endoreplication in response to conditional stresses and environmental inputs, for example during tissue regeneration and cancer (Fox and Duronio, 2013; Losick et al., 2013; Øvrebø and Edgar, 2018). It is still not fully understood, however, how remodeling of the cell cycle promotes the switch from mitotic cycles to endoreplication cycles in development and disease.

There are common themes across plants and animals for how cells switch to endoreplication during development. One common theme is that developmental signaling pathways induce endoreplication by inhibiting the mitotic cyclin dependent kinase 1 (CDK1). After CDK1 activity is repressed, repeated G / S cell cycle phases are controlled by alternating activity of the ubiquitin ligase APC/C^CDH1^ and Cyclin E / CDK2 (Edgar et al., 2014). Work in *Drosophila* has defined mechanisms by which APC/C^CDH1^ and CycE /Cdk2 regulate G / S progression and ensure that the genome is duplicated only once per cycle (Edgar et al., 2014; Hong et al., 2007; Lilly and Spradling, 1996; Narbonne-Reveau et al., 2008; Zielke et al., 2011; Zielke et al., 2008). Despite these conserved themes, how endoreplication is regulated can vary among organisms and tissues within an organism. These variations include the identity of the signaling pathways that induce endoreplication, the mechanism of CDK1 inhibition, and the downstream effects of cell cycle remodeling into either an endomitotic cycle (partial mitosis) or endocycle (skip mitosis) (Edgar et al., 2014; Fox and Duronio, 2013). In many cases, however, the identity of the developmental signals and the molecular mechanisms of cell cycle remodeling are not known.

We had previously used two-color microarrays to compare the transcriptomes of endocycling and mitotic cycling cells in *Drosophila* tissues (Maqbool et al., 2010). We found that endocycling cells of larval fat body and salivary gland have dampened expression of genes that are normally induced by E2F1, a surprising result for these highly polyploid cells given that many of these genes are required for DNA synthesis. Nonetheless, subsequent studies showed that this property is conserved in endoreplicating cells of mouse liver, megakaryocytes, and trophoblast giant cells (Chen et al., 2012; Pandit et al., 2012; Zielke et al., 2011). *Drosophila* endocycling cells also had dampened expression of genes regulated by the Myb transcription factor, the ortholog of the human B-Myb oncogene (Katzen et al., 1998; Maqbool et al., 2010). Myb is part of a larger complex called Myb-MuvB (MMB), whose subunit composition and function are conserved from flies to humans (Beall et al., 2002; Guiley et al., 2018; Korenjak et al., 2004; Lewis et al., 2004; Sadasivam et al., 2012). A major conserved function of the MMB is promoting periodic transcription of genes that are required for mitosis and cytokinesis (Blanchard et al., 2014; DeBruhl et al., 2013; Georlette et al., 2007; Guiley et al., 2018; Wen et al., 2008). It was these mitotic and cytokinetic genes whose expression was dampened in *Drosophila* endocycles, suggesting that this repressed Myb transcriptome promotes the switch to endocycles that skip mitosis. It is not known, however, how E2F1 and Myb activity are downregulated, nor which downstream genes are key for the remodeling of mitotic cycles into endocycles.

In addition to endoreplication during development, there are a growing number of examples of cells switching to endoreplication cycles in response to conditional stresses and environmental inputs (Fox and Duronio, 2013; Øvrebø and Edgar, 2018). We will call these induced endoreplicating cells (iECs) to distinguish them from developmental endoreplicating cells (devECs). For example, iECs contribute to tissue regeneration after injury in flies, mice, humans, and the zebrafish heart, and evidence suggests that a transient switch to endoreplication contributes to genome instability in cancer (Cao et al., 2017; Chen et al., 2016; Cohen et al., 2018; González-Rosa et al., 2018; Herget et al., 1997; Losick et al., 2013; Losick et al., 2016). Hypertension also promotes an endoreplication that increases the size and ploidy of heart muscle cells, and this hypertrophy contributes to cardiac disease (Dominiczak et al., 1996; Herget et al., 1997). Experimental inhibition of CDK1 activity is also sufficient to induce endoreplication in flies, mouse, and human cells (Davoli and de Lange, 2011; Diril et al., 2012; Hassel et al., 2014; Hayashi, 1996; Itzhaki et al., 1997; Sauer et al., 1995). We previously showed that these experimental iECs in *Drosophila* are similar to devECs in that they skip mitosis, have oscillating CycE / Cdk2 activity, periodically duplicate their genome, and repress the apoptotic response to genotoxic stress (Hassel et al., 2014; Maqbool et al., 2010; Qi and Calvi, 2016; Zhang et al., 2014). In general, however, it is little understood how similar the cell cycles of iECs are to devECs. In this study, we use experimental iECs in *Drosophila* to investigate how the cell cycle is remodeled when cells switch from mitotic cycles to endoreplication cycles, and how similar this remodeling is between iECs and devECs.

## Results

### Induced endocycling cells have reduced expression of Myb-regulated, but not E2F1-regulated, genes

We sought to understand how remodeling of the cell cycle program determines the switch from mitotic cycles to endoreplication cycles, and how similar iECs are to devECs. As a model for iECs, we induced *Drosophila* S2 cells in culture into endoreplication cycles by treating them with Cyclin A (CycA) double-stranded RNA (dsRNA), which we and others had previously shown induces endocycles (Hassel et al., 2014; Mihaylov et al., 2002; Sauer et al., 1995). As a control, we analyzed mitotic cycling S2 cells that were treated in parallel with dsRNA for GFP, permitting a comparison of variant cell cycles in the same cell type. Flow profiling 96 hours after treatment with CycA dsRNA indicated that more than 50% of cells had a polyploid DNA content of ≥ 8C, and a commensurate reduced fraction of cells with diploid 2C and 4C contents (Figure 1A, B). Importantly, these cells had genome doublings of 8C, 16C, and 32C that were multiples of the diploid DNA content, suggesting that they duplicate their genome through repeated G / S endocycles (Figure 1A,B). In contrast, knockdown of the mitotic Cyclin B (CycB) did not induce cells to endoreplicate, perhaps because of functional redundancy with CycB3 (Jacobs et al., 1998; McCleland et al., 2009) (Figure S1). These results confirm previous findings that knockdown of CycA is sufficient to induce endoreplication in S2 cells (hereafter CycA dsRNA iEC) (Hassel et al., 2014).

**Figure 1.**
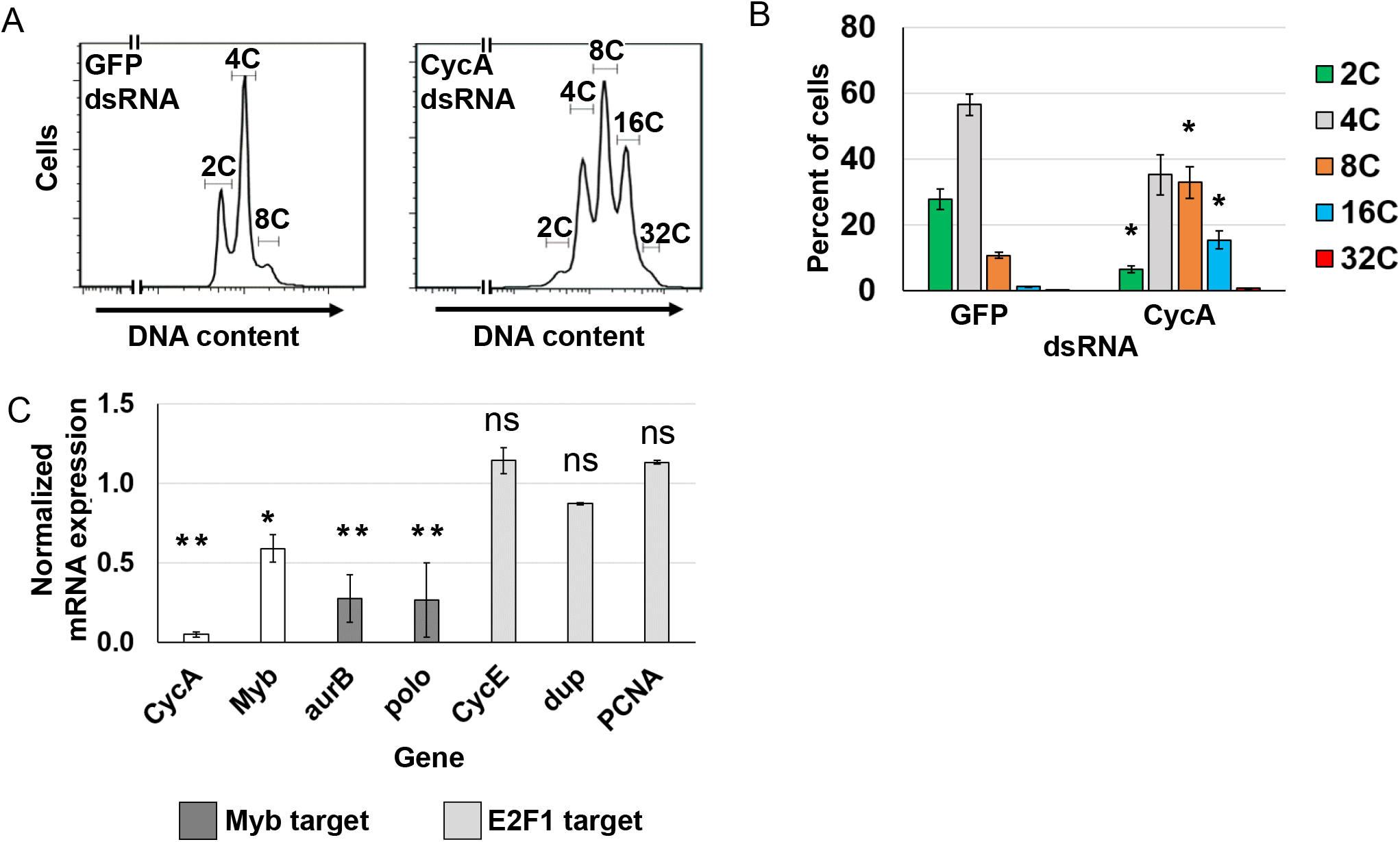
Knockdown of Cyclin A induces endoreplication cycles. **(A)** Flow cytometry of DNA content in propidium iodide labeled S2 cells 96 hours after treatment with either GFP dsRNA (control) or CycA dsRNA. **(B**) Quantification of the ploidy classes after GFP or CycA dsRNA treatment. Mean and S.E.M. for N=3, * - p <0.05 comparing each CycA dsRNA ploidy class with the corresponding ploidy class in control GFP dsRNA treated cells. **(C)** qRT-PCR analysis of select Myb and E2F1 target gene expression in CycA dsRNA iECs. Normalized mRNA is the ratio of mRNA levels in CycA dsRNA divided by control GFP dsRNA cells (N=3, mean and S.E.M., * - p < 0.05, ** - p < 0.01, ns – not significant).

We had previously shown that endocycling cells (G/S cycle) of the *Drosophila* larval salivary gland and fat body have dampened expression of genes that are normally induced by E2F1 and the MMB transcription factors (Maqbool et al., 2010). To determine if this change in transcriptome signature also occurs in CycA dsRNA iECs, we analyzed the expression of several candidate genes whose expression is induced by E2F1 or MMB. RT-qPCR results indicated that CycA dsRNA iECs had reduced expression of the Myb subunit of the MMB and two genes that are positively regulated by the MMB and essential for mitosis (*aurora B* and *polo*) (Figure 1C). In contrast, the expression of three genes normally induced by E2F1 at G1 / S (*Cyclin E, double-parked*, and *PCNA*) were similar between CycA dsRNA iECs and mitotic cycling cells (Figure 1C). These results suggest that CycA dsRNA iECs are similar to developmental endocycling cells (devECs) in that they have reduced expression of MMB-target genes, but differ in that they do not have reduced expression of genes that require E2F1 for their expression.

### Knockdown of *Cyclin A* or *Myb* induces similar endoreplication cycles

Although CycA dsRNA iECs had lower expression of two MMB-induced genes that are required for mitosis, it was unclear whether dampened MMB activity contributed to the switch to endoreplication. To address this question, we knocked down expression of the MMB Myb subunit, which is required to induce the expression of genes for mitosis and cytokinesis (Blanchard et al., 2014; DeBruhl et al., 2013; Georlette et al., 2007). Knockdown of Myb inhibited cell proliferation, and resulted in an increase in polyploid DNA content that was similar to that of CycA dsRNA iECs (Figure 2A, B, Figure S2). We then used fluorescent microscopy to further evaluate ploidy and cell cycle in CycA and Myb knockdown cells. Cells were incubated in the nucleotide analog EdU for two hours followed by fluorescent click-it labeling of EdU to detect cells in S phase, labeled with antibodies against phospho-histone H3 (PH3) to detect chromosomes in mitosis, and incubated with DAPI to label nuclear DNA (Goto et al., 1999; Hendzel et al., 1997; Hooser et al., 1998). Imaging of DAPI-labeled nuclei showed that treatment of cells with either CycA or Myb dsRNA resulted in a similar frequency and size of large polyploid nuclei, indicating that Myb knockdown induced endoreplication (hereafter Myb dsRNA iEC) (Figure 2C-F). There was a higher fraction of multinucleate Myb dsRNA iECs (~15%) than CycA dsRNA iECs (~8%), suggesting that Myb knockdown results in a somewhat larger fraction of endomitotic cells than does CycA knockdown (Figure 2G). Approximately 30% of CycA dsRNA iECs and Myb dsRNA iECs incorporated EdU, a frequency that was similar in both mononucleate and multinucleate populations, consistent with periodic duplications of the genome during both endocycles and endomitotic cycles (Figure 2H). Despite this evidence for periodic endoreplication, the fraction of total cells with mitotic PH3 labeling was not decreased after CycA knockdown (~5%), and was slightly increased after Myb knockdown in the mononucleate population (~10%) (Figure 2H). Unlike control mitotic cells, however, the PH3 labeling after CycA and Myb knockdown was diffuse, with little evidence of fully condensed mitotic chromosomes, suggesting that these cells were either arrested or delayed in early mitosis or endomitosis (Figure 2C-E’). These results indicate that knockdown of Myb is sufficient to induce endoreplication cycles that are similar to those after knockdown of CycA.

**Figure 2.**
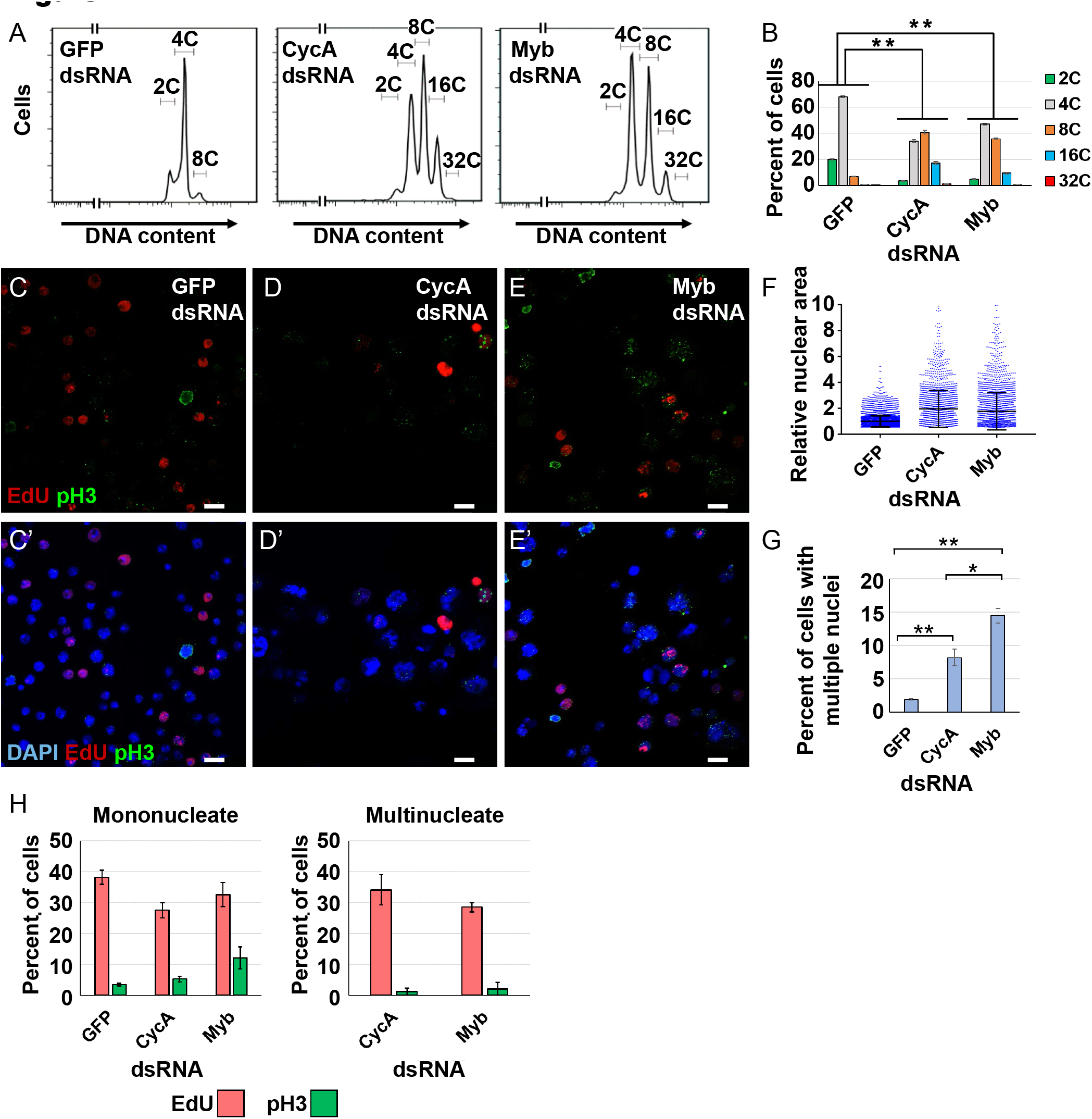
Myb or CycA knockdown induce similar endoreplication cycles. **(A)** Flow cytometry of DNA content in S2 cells treated with the indicated dsRNAs for 96 hours. **(B)** Quantification of the induced polyploidy after Myb dsRNA or CycA dsRNA treatment. Mean and S.E.M., N=3, ** - p <0.01 for each ploidy class compared to GFP dsRNA. (**C-E’**) Micrographs of cells labeled with EdU (red), pH3 (green) and DAPI (blue in **C’-E’**) after four days of treatment with dsRNA for GFP (control) **(C,C’)**, CycA **(D,D’)**, or Myb **(E,E’)**. Scale bars are 10μM. **(F)** Knockdown of CycA or Myb increases nuclear size. Quantification of nuclear area of S2 cells after knockdown of GFP, CycA or Myb. Each dot represents the nuclear area of a single cell divided by the mean area of GFP controls (machine units) (Mean and S.D. N=3). **(G)** Quantification of the fraction of total S2 cells with multiple nuclei after the indicated treatment (mean and S.E.M. for N=3, * - p<0.05, ** - p<0.01 relative to GFP dsRNA). **(H)** Quantification of EdU and pH3 labeling in mononucleate and multinucleate cells after treatment with the indicated dsRNAs (mean and S.E.M. for N=3).

### Myb induction of M phase gene expression is downstream of and dependent on CycA

The similarity between CycA dsRNA and Myb dsRNA iECs suggested that they may have a functional relationship. To evaluate this, we examined cell cycle protein levels by Western blotting. This analysis showed that CycA and Myb dsRNA treatments resulted in lower levels of the respective proteins (Figure 3A). Myb dsRNA iECs also had greatly reduced levels of CycB protein, which is consistent with previous evidence that the MMB induces transcription of the *CycB* gene during mitotic cycles (Figure 3A) (Georlette et al., 2007; Herget et al., 1997; Okada et al., 2002). CycA dsRNA iECs also had lower CycB protein, suggesting that CycA knockdown may compromise MMB activity, consistent with the RT-qPCR results that two other Myb target genes, *aurB* and *polo*, are expressed at lower levels in CycA dsRNA iECs (Figure 1C). We extended this RT-qPCR analysis and measured the mRNA levels for eight known MMB target genes that function in mitosis or cytokinesis. Knockdown of either CycA or Myb reduced the expression of all these genes to similar extents (Figure 3B). Knockdown of CycA resulted in reduced Myb mRNA, whereas knockdown of Myb did not reduce levels of CycA mRNA, suggesting that Myb knockdown is sufficient to induce endoreplication cycles even when CycA levels are not reduced, consistent with the Western results (Figure 3A, B). These results suggest that CycA is required for MMB transcriptional activation of M phase genes.

**Figure 3.**
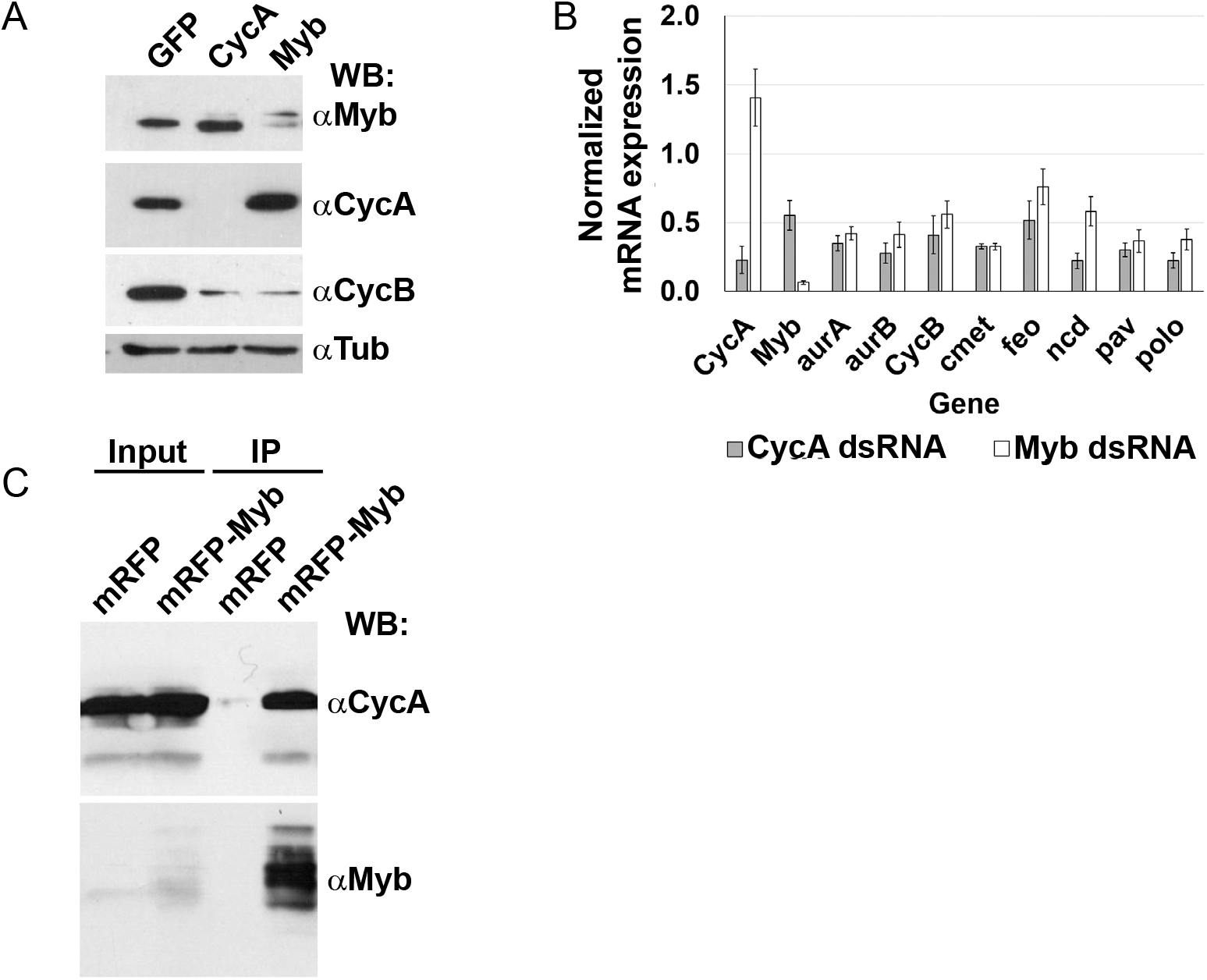
Myb induction of M phase gene expression is downstream of and dependent on CycA. **(A)** Cyclin B protein levels are reduced in CycA^dsRNA^ and Myb^dsRNA^ iECs. Western blot of S2 cell extracts after treatment with the indicated dsRNA and incubated with antibodies against CycA, Myb, CycB, and alpha-Tubulin (loading control) (N=3,a representative blot is shown). **(B)** qRT-PCR analysis of select MMB target gene expression in CycA^dsRNA^ and Myb^dsRNA^ iECs. Values shown are the average ratio of mRNA levels versus control GFP^dsRNA^ cells (mean and S.E.M. for N=3 biological replicates). **(C)** CycA and Myb proteins interact *in vivo*. Larvae expressing *UAS-CycA* and either *UAS-mRFP* or *UAS-mRFP-Myb* were immunoprecipitated with nanobodies against mRFP and then Western blotted (WB) with antibodies against CycA or Myb (representative blot, N=3).

To evaluate if CycA regulation of the MMB could be direct, we determined if Myb and CycA physically interact. We used the GAL4 / UAS system to express *UAS-CycA* with either *UAS-Myb-RFP* or *UAS-RFP* in mitotic cycling imaginal discs, immunoprecipitated Myb-RFP or RFP with an anti-RFP nanobody, and then blotted for Cyclin A (Weiss et al., 1998; Wen et al., 2008). The results indicated that Myb-RFP, but not RFP alone, co-precipitates with CycA (Figure 3C). All together, these results suggest that during mitotic cycles a CycA / CDK complex is directly required for MMB to induce expression of genes required for M phase, and that in the absence of this activation cells switch to endoreplication cycles.

### CycA dsRNA and Myb dsRNA iECs have reduced expression of Myb-regulated genes that function at multiple steps of mitosis and cytokinesis

The results were consistent with the model that during the mitotic cell cycle CycA / Cdk1 activates the MMB to induce the expression of genes required for mitotic entry and progression. To further evaluate the relationship between CycA and Myb and gain insight into remodeling of mitotic cycles into endoreplication cycles, we analyzed the global transcriptomes of CycA dsRNA and Myb dsRNA iECs by RNA-Seq. The transcriptome of these two iEC populations were compared to control mitotic cycling cells treated in parallel with GFP dsRNA, all in three biological replicates. Genes were defined as differentially expressed (DE) in iECs if their normalized steady state mRNA levels differed from mitotic cycling cells with a log2 fold change (log2FC) of at least +/- 0.5 and a false discovery rate (FDR) corrected q-value <0.05 (Benjamini and Hochberg, 1995).

The RNA-Seq results indicated that a switch from mitotic cycles to endoreplication in CycA dsRNA and Myb dsRNA iECs is associated with differential expression of thousands of genes (Figure 3A, Tables S1 and S2). Comparison of the CycA dsRNA and Myb dsRNA iEC transcriptomes revealed that they shared a total of 966 genes that were differentially expressed compared to mitotic cycling controls (698 increased and 268 decreased) (Figure 4B, Tables S3 and S4). Gene Ontology (GO) analysis indicated that the shared upregulated genes were significantly enriched for genes involved in immunity, metabolism, and development, and that shared down regulated genes also included those for energy metabolism (Figure S3, S4, Table S3, S4). Relevant to cell cycle remodeling, CycA dsRNA and Myb dsRNA iECs both had significantly lower expression of 47 genes that are required for different processes in mitosis (Figure S4, Table S4). It has been previously shown that the MMB is required for expression of these genes during G2 and early M phase, and that many are direct targets of the MMB in Drosophila Kc cells (Georlette et al., 2007). These results further suggest that CycA is required to activate transcriptional induction by the MMB, and that downregulation of this MMB transcriptome in CycA dsRNA and Myb dsRNA iECs may contribute to the switch from mitotic cycles to endoreplication cycles.

**Figure 4.**
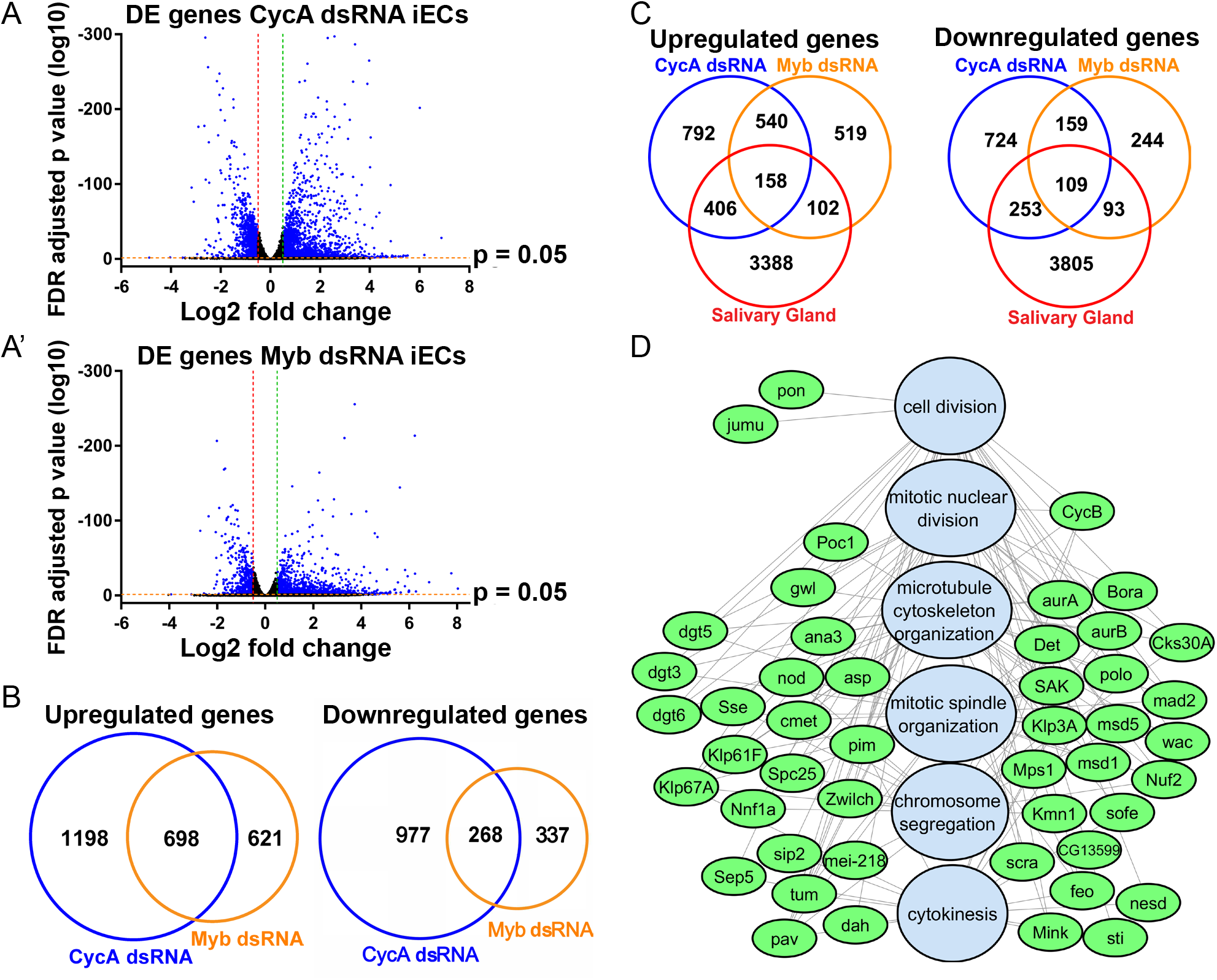
Both iECs and devECs have reduced expression of Myb target genes that function at multiple steps of mitosis and cytokinesis. **(A, A’)** Volcano plots of RNA-Seq results for differentially expressed (DE) genes in CycA dsRNA iECs **(A)** and Myb dsRNA iECs **(A’)** each relative to GFP dsRNA control cells (N=3 biological replicates). Vertical red and green dotted lines indicate thresholds for log2 fold change (< −0.5 and >+0.5) in iECs and horizontal red line the FDR adjusted p-value (<0.05). Blue dots represent genes that fulfill both of these criteria. **(B)** Venn diagrams comparing the overlap of DE genes in CycA dsRNA and Myb dsRNA relative to control GFP dsRNA cells. See also Tables S1-S3. **(C)** Venn diagrams comparing the overlap of DE genes in iECs with DE genes in salivary glands (SG) (relative to mitotic brains and discs). **(D)** Gene Ontology (GO) analysis of genes downregulated in iECs and devECs indicate an enrichment for Myb target genes that are required for mitosis. Shown is a network analysis with GO biological process categories in blue and downregulated genes in green. See also Table S4, S5.

We wished to use the RNA-Seq results to address how similar iECs are to devECs. Our previous transcriptome analysis of endocycling cells of *Drosophila* tissues used two-color expression microarrays and represented a limited gene set (Maqbool et al., 2010). We therefore repeated the analysis with RNA-Seq, comparing the transcriptome of endocycling larval salivary glands (SG) to that of mitotic cycling brains and discs (B-D), in three biological replicates. The RNA-Seq results were consistent with our previous array analysis, and showed that expression of both the E2F1 (mostly S phase genes) and MMB transcriptomes (mostly M phase genes) are dampened in endocycling SG cells relative to mitotic cycling B-D cells. A comparison with iECs showed that devECs and both types of iECs have in common 158 genes that are increased and 109 genes that are decreased in expression relative to mitotic cycling cells (Figure 4C). While devECs have a dampened E2F1 transcriptome, iECs did not, consistent with our RT-qPCR results (Figure 1C). However, among the shared downregulated genes the most significantly enriched GO categories included the 47 MMB-induced genes that have multiple functions during mitosis and cytokinesis (Figure 4D, Figure S5 Table S5). These genomic results show that iECs and devECs share the property of having a dampened MMB transcriptome of genes that function in mitosis, but differ in that only devECs have a dampened E2F1 transcriptome.

### Integration of genetic analysis with RNA-Seq implicates a CycA – Myb - AurB network in endoreplication control

The findings in S2 cells suggested that CycA / Cdk1 activates the MMB to induce transcription of a battery of genes required for mitosis, and that knockdown of CycA or Myb results in reduced mitotic gene expression, thereby promoting endoreplication cycles. It was unclear, however, which of the downregulated mitotic genes downstream of Myb are key for the decision to switch to endoreplication cycles. To address this, we took an integrative genetic approach, using a collection of fly strains with GAL4-inducible UAS-dsRNAs to knock down the expression of genes that RNA-Seq showed were downregulated. We used an inclusive criterion and knocked down genes that were downregulated by log2 fold of at least −0.5 in both types of iECs, but without regard to p value (n=242 available strains, n=240 genes) (Ni et al., 2011) (Table S6). *dpp-GAL4* was used to express these dsRNAs along the anterior-posterior compartment boundary of the larval wing disc, and then the hair pattern of adult wings was examined (Staehling-Hampton et al., 1994) (Figure 5A). Each hair on the adult wing represents an actin protrusion from a single cell, and it is known that polyploidization of wing cells results in fewer and larger hairs (Guild et al., 2005; Hanson et al., 2005; Mitchell et al., 1983) (Figure 5A). As proof of principle, expression of a *UAS-CycA^dsRNA^* along the A/P boundary resulted in a central stripe of longer hairs on the adult wing surface and margin between veins L3 and L4, with many cells producing clusters of multiple hairs (Figure 5B). Knockdown of Myb also resulted in a stripe of larger and more widely spaced wing hairs (Figure 5C). Although the Myb knockdown phenotype was less severe than that of CycA knockdown, this Myb dsRNA is inefficient and only knocks down Myb mRNA to ~50% of wild type levels (DNS). Knockdown of either CycA or Myb resulted in reduced spacing between wing veins L3 and L4, suggesting that a switch to polyploid cycles and increase in cell size was not able to completely recapitulate normal tissue growth (Figure 5B, C). Among the 242 strains tested, 23 resulted in lethality before adulthood, suggesting that their functions are essential, but were otherwise uninformative (Table S6).

**Figure 5.**
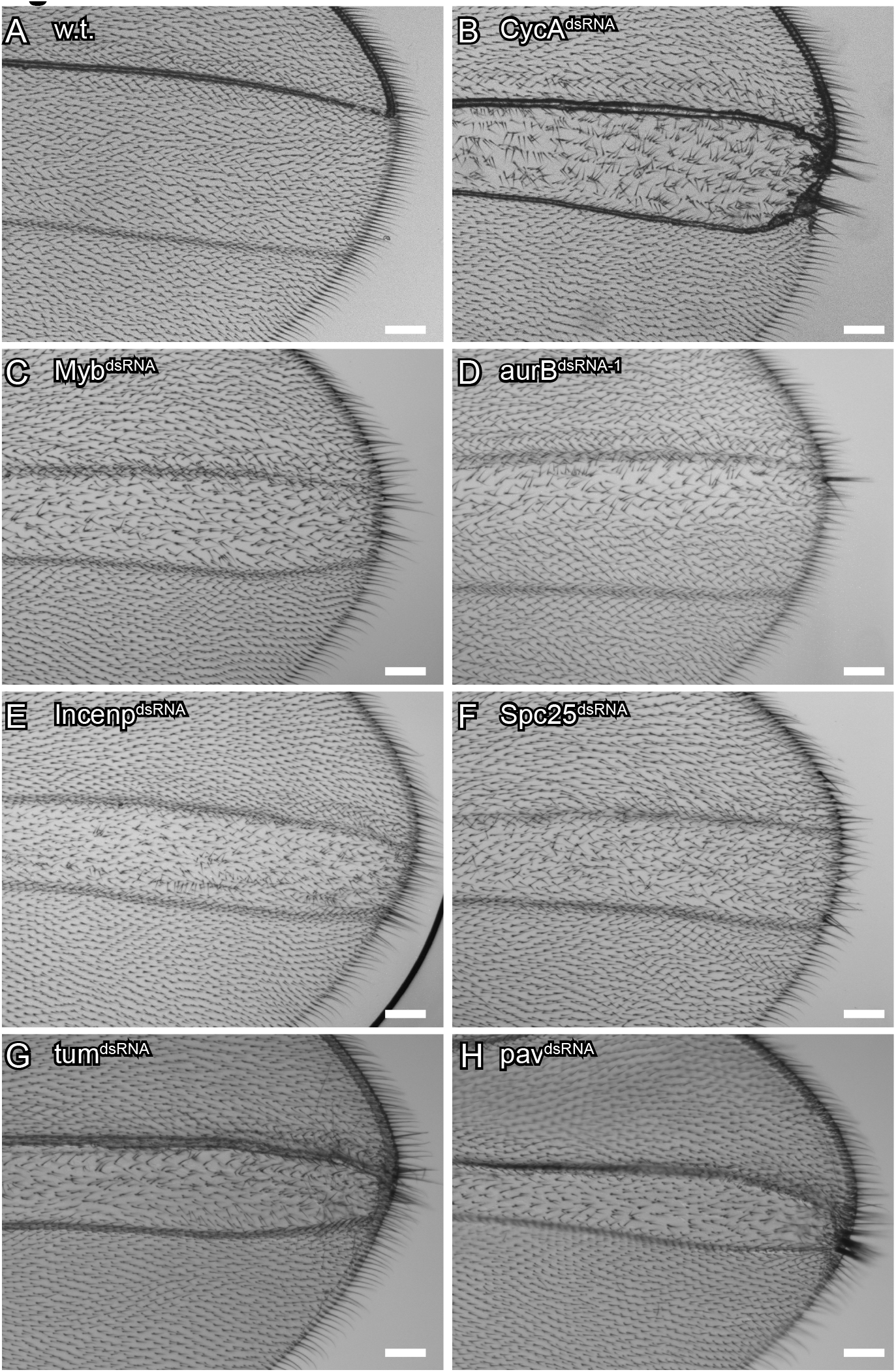
Integration of a genetic screen with RNA-Seq implicates a CycA-Myb-AurB network in endoreplication control. Positive results from an RNAi screen of candidate genes that were expressed at lower levels in iECs. Shown are brightfield images of an adult wing from animals that expressed *dpp-GAL4* and the indicated UAS-dsRNA along the A-P margin of larval wing discs. **(A)** A control wild type (w.t.) wing from a *dpp:GAL4; UAS-GFP* animal. **(B)** A wing from a *dpp-GAL4; UAS-CycA^dsRNA^* animal. Note clusters of larger and thicker hairs on the wing surface along the A-P boundary as well as on the distal wing margin between veins L3 and L4. **(C-H)** Adult wings after *dpp-GAL4* induced expression of *UAS-Myb^dsRNA^* **(C)**, *AurB^dsRNA-1^* (**D**), *Incenp^dsRNA^* **(E)**, *Spc25^dsRNA^* **(F)**, *tum^dsRNA^* **(G)**, or *pav^dsRNA^* **(H)**. Anterior is up, Scale bar is 15μm.

Among the other 217 crosses that survived to adults, knockdown of five genes reproducibly resulted in enlarged wing hairs, *aurora B* (*aurB*), *Incenp, Spc25, tumbleweed (tum), and pavarroti (pav)* (Figure 5D-F) (Adams et al., 2001; Adams et al., 1998; Giet and Glover, 2001; Goshima et al., 2007; Somers and Saint, 2003). Remarkably, all of these genes are either part of the chromosomal passenger complex (CPC) or are downstream effectors of it. AurB kinase and INCENP are two subunits of the four-subunit CPC complex, which phosphorylates downstream targets to regulate multiple processes of mitosis and cytokinesis (Carmena et al., 2015; Carmena et al., 2012). Spc25 is a subunit of the Ndc80 outer kinetochore complex, which is phosphorylated by the CPC to regulate microtubule-kinetochore attachments (Nannas and Murray, 2012; Umbreit et al., 2012). The CPC also phosphorylates the Tum protein, a Rac-GAP protein that regulates the kinesin Pav for proper cytokinesis (Tao et al., 2016). While knockdown of any of these five genes resulted in longer hairs on the wing margin, knockdown of *aurB* had the strongest effect on hair length in the anterior half of the L3 / L4 intervein region (Figure 5 D-H). These genetic results, together with the evidence that these genes are repressed in iECs, suggest that a dampened CycA – Myb – AurB network promotes a switch from mitotic cycles to endoreplication cycles.

### Knockdown of a CycA – Myb – AurB network induces endoreplication in ovarian follicle cells

The combined RNA-Seq and genetic results suggested that the status of a CycA – Myb – AurB network determines the decision between mitotic and endoreplication cycles. To quantify the effects of this network on cell cycles, we conducted a genetic mosaic analysis in somatic follicle cells of the ovary, which have several methodological advantages. Follicle cells form a regular epithelial sheet around 15 germline nurse cells and one oocyte in each maturing egg chamber. Their cell cycle programs are well characterized and coupled with stages of oogenesis, dividing mitotically during stages 1-6, undergoing three endocycles during stages 7-10A, and then selectively re-replicating genes required for eggshell synthesis during stages 10B-14 (Calvi et al., 1998; Deng et al., 2001; Mahowald, 1972). We had shown previously that knockdown of *CycA* or over-expression of *Fzr* (Cdh1) was sufficient to induce mitotic follicle cells into a precocious endocycle before stage 7 (Hassel et al., 2014; Qi and Calvi, 2016). To achieve conditional knockdown of the genes identified in the wing screen, we used the heat-inducible GAL4 / UAS FLP-On system, which results in clonal activation of GAL4 and induction of a *UAS-dsRNA* and a *UAS-RFP* reporter in a subset of cells, (Pignoni and Zipursky, 1997). Three days after heat induction, we quantified the number of cells in the clone, their nuclear size, and their DNA content by measuring DAPI fluorescence. Cell number was compared to control wild type clones, and nuclear size and DNA content was normalized to wild type follicle cells outside of the clone in the same egg chamber. If a gene knockdown induces endoreplication, it will result in clones with fewer cells that have an increase in nuclear size and DNA content.

We used confocal imaging to analyze clones in stage 6, the latest stage of oogenesis during which follicle cells mitotically divide. Wild type, control clones were comprised of ~28 RFP-positive cells, indicating that they had divided ~4-5 times since FLP-On in the original single founder cell three days earlier in oogenesis (Figure 6A, A’, K, L). FLP-On of *UAS-CycA^dsRNA^* resulted in clones with only one to three cells, each with a single large nucleus that had an increased DNA content up to ~16C (Figure 6B, B’, K, L, Table I). This result indicated that they had switched to endocycles during the first or second mitotic cycle after *CycA* knockdown, consistent with our previously published results (Hassel et al., 2014). Expression of the *UAS-Myb^dsRNA^* resulted in some clones with reduced cell numbers and larger nuclei, suggesting that some cells had switched to endoreplication, but with variable expressivity among clones (Figure 6C, C’, K, L, Table I). This variably expressive phenotype is likely the result of partial Myb knockdown by the *UAS-Myb^dsRNA^* transgene. A few Myb knockdown cells had two nuclei that were increased in size and DNA content, with some labeled by EdU, suggesting that these cells had failed cytokinesis before replicating their DNA again, a type of endomitosis (DNS). These results are consistent with the results in S2 cells that indicate that knockdown of CycA or Myb is sufficient to induce endoreplication.

**Table I.**
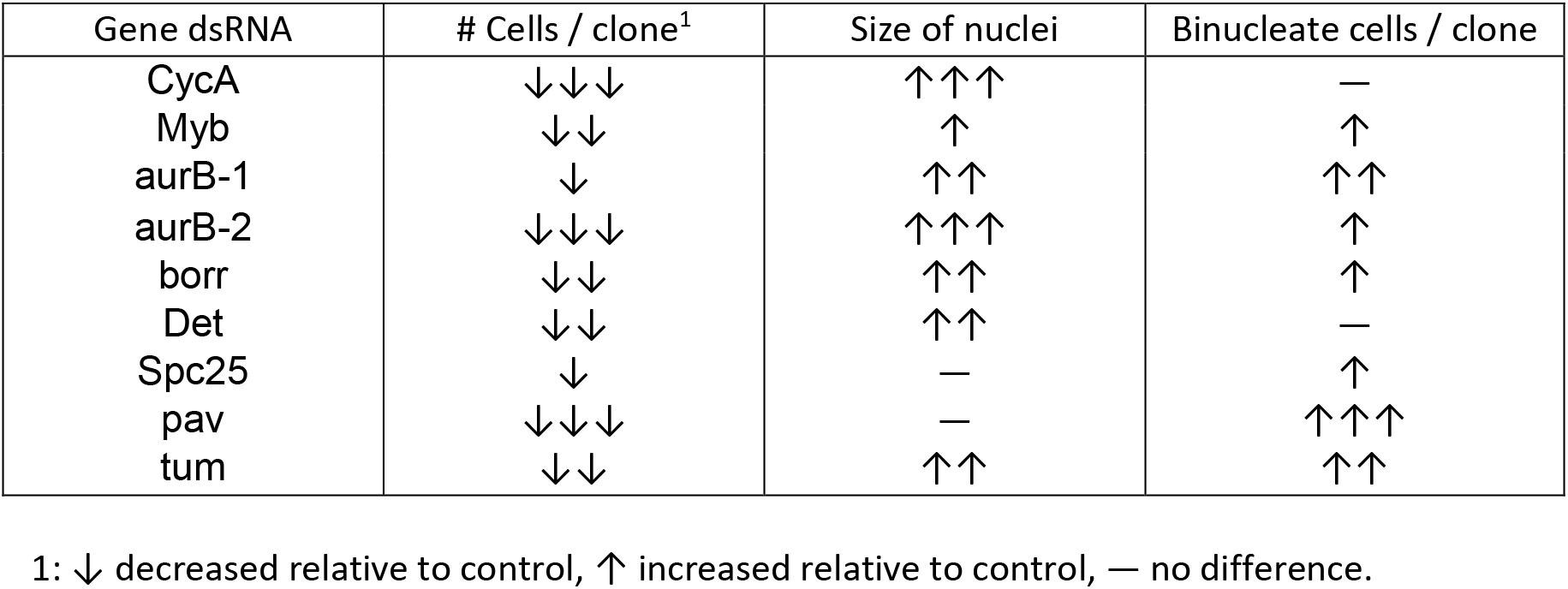
Summary of follicle cell clone phenotypes.

| Gene dsRNA | # Cells / clone<sup>1</sup> | Size of nuclei | Binucleate cells / clone |
| --- | --- | --- | --- |
| CycA | ↓↓↓ | ↑↑↑ | — |
| Myb | ↓↓ | ↑ | ↑ |
| aurB-1 | ↓ | ↑↑ | ↑↑ |
| aurB-2 | ↓↓↓ | ↑↑↑ | ↑ |
| borr | ↓↓ | ↑↑ | ↑ |
| Det | ↓↓ | ↑↑ | — |
| Spc25 | ↓ | — | ↑ |
| pav | ↓↓↓ | — | ↑↑↑ |
| tum | ↓↓ | ↑↑ | ↑↑ |

1: ↓ decreased relative to control, ↑ increased relative to control, — no difference.

**Figure 6.**
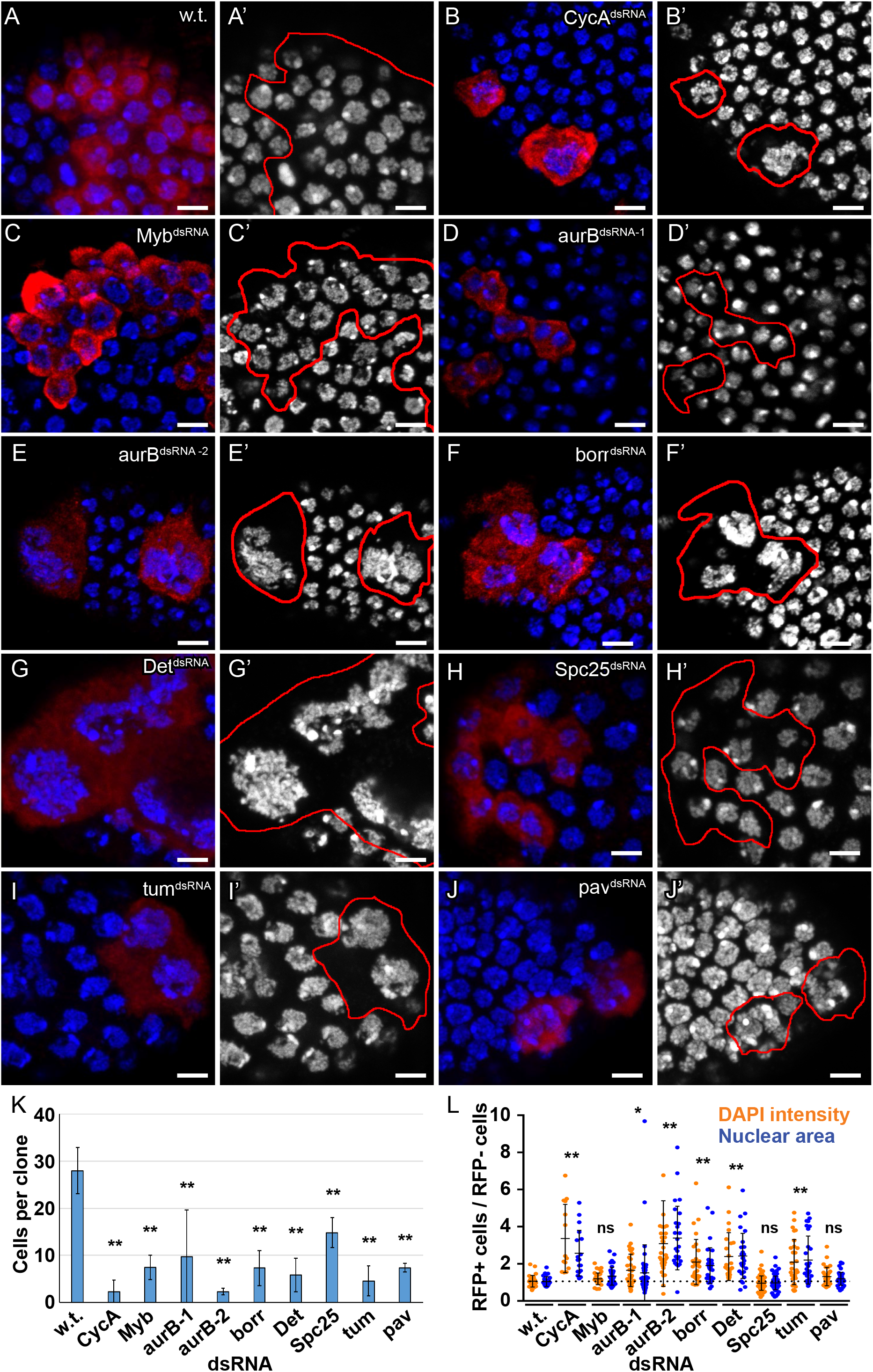
Knockdown of a CycA – Myb – AurB network induces endoreplication in ovarian follicle cells. **(A-J’)** Clonal knockdown in follicle cells. Clonal FLP-On of GAL4 expression was induced by heat treatment of adult females of genotype *hsp70-FLP, Act>cd2>Gal4, UAS-mRFP* and the indicated *UAS-dsRNA*. Clones in stage 6 egg chambers were analyzed three days later by labeling with anti-mRFP (red) and DAPI (blue in **A-J** and white in **A’-J’**). Red outlines in **A’-J’** indicate clone boundaries. **(K)** Quantification of cells per clone. RFP labeled cells were counted from at least four independent clones. More clones were counted in samples with fewer cells / clone (mean and S.D. for N ≥ 4 clones, ** - p<0.01 relative to wild type control clones). **(L)** Quantification of nuclear area and DAPI intensity. The nuclear area (blue dots) and DAPI intensity (orange dots) of single cells in a clone (RFP+) were measured and divided by the mean nuclear area and DAPI intensity of wild type cells (RFP-) in the same egg chamber (mean and S.D. for N ≥ 3 clones, * - p<0.05, ** - p<0.01, ns – not significant relative to wild type, control cells).

The combined RNA-seq and genetic screen results suggested that reduced expression of CPC subunits downstream of Myb contributes to the switch to endoreplication. The expression of *UAS-AurB^dsRNA-1^* in wing discs had resulted in elongated hairs, and in follicle cells resulted in clones of only two to three cells, with variable increases in nuclear size and DNA content (Figure 6D, D’ K, L, Table I). A few of these cells had two nuclei of increased size and DNA content that labeled with EdU, suggesting that *UAS-AurB^dsRNA-1^* impaired cytokinesis followed by endoreplication in a subset of cells (DNS). This *UAS-AurB^dsRNA-1^* transgene is based on a series of vectors that are not highly efficient for dsRNA expression. Expression of a more efficient *UAS-AurB^dsRNA-2^* had resulted in lethality before adulthood in the wing screen (Table S6). FLP-ON expression of this *UAS-AurB^dsRNA-2^* in follicle cells resulted in clones composed of only one to two cells, each with a single, large, polyploid nucleus (Figure 6E, E’, K, L). Many of these nuclei were multi-lobed, with connected chromatin masses composed of large chromosomes that appeared polytene (Figure 6E, E’). These results suggest that mild knockdown of AurB results in cytokinesis failure, whereas a more severe knockdown results in a failure to segregate chromosomes followed by endoreplication.

To further test whether reduced CPC activity induces endoreplication, we knocked down expression of its other three subunits Incenp, Borealin-related (Borr), and Deterin (Det) (fly Survivin ortholog), all of which were expressed at lower levels in iECs and salivary glands (Tables S4, S5) (Adams et al., 2001; Eggert et al., 2004; Jones et al., 2000). Although expression of *UAS-Incenp^dsRNA^* in the wing resulted in a mild phenotype, in follicle cells it did not reduce the number of cells per clone nor increase DNA content, an uninformative result because this UAS transgene is optimized for expression in the germline but expressed poorly in the soma (Table I, and DNS). In contrast, the use of optimized dsRNA transgenes to knockdown the other subunits of the CPC, Borr and Det, resulted in clones with very few cells with large, polyploid nuclei (Figure 6 F-G’, K, L, Table I). These results further indicate that reduction of CPC activity is sufficient to switch cells from mitotic cycles to endoreplication cycles.

The CPC has multiple functions during mitosis and cytokinesis, all of which are impaired by knockdown of the CPC subunits (Carmena et al., 2012). Three genes that encode proteins that function downstream of the CPC were expressed at lower levels in iECs and recovered in the wing screen – *Spc25, tum, and pav*. Spc25 mediates CPC regulation of microtubule-kinetochore attachments (Janke et al., 2001; Williams et al., 2007). Spc25 knockdown in follicle cells resulted in fewer cells per clone, but the nuclear size and DNA content of these cells were not increased, indicating that knockdown of Spc25 inhibits cell division but is not sufficient to induce endoreplication (Figure 6H,H’,K, L, Table I). The Rac-GAP Tum is phosphorylated by the CPC, and promotes cytokinesis in part through its interaction with the kinesin Pav (Adams et al., 1998; Tao et al., 2016). Knockdown of either Tum or Pav in follicle cells resulted in fewer cells per clone, with many binucleate, indicating that cytokinesis was inhibited (Figure 6 I-K, Table I). EdU labeling indicated that some of these binucleate cells entered S phase after cytokinesis failure, with Tum knockdown cells having an increase in nuclear size and DNA content (mean ~2 fold, max ~4 fold increase) (Figure 6I, I’, K, L and DNS). These results suggest that inhibition of cytokinetic functions downstream of the CPC is sufficient to induce an endoreplication cycle. All together, these results suggest that the status of a CycA – Myb – AurB network determines the choice between mitotic and endoreplication cycles.

## Discussion

We have investigated how the cell cycle is remodeled when mitotic cycling cells are induced to switch into endoreplication cycles. We have found that repression of a CycA – Myb – AurB mitotic network induces a switch to endoreplication. This network was also downregulated in developmental endocycles of the larval salivary gland, revealing a similarity in the regulation of iEC and devEC cell cycles. Unlike devECs, however, iECs did not have reduced expression of E2F1 induced genes, uncovering a diversity of cell cycle transcriptome remodeling during endoreplication. Overall, these findings define how cells either commit to mitosis or switch to different types of endoreplication cycles, with broader relevance to understanding the regulation of these variant cell cycles and their contribution to development, tissue regeneration, and cancer.

Our findings indicate that the status of the CycA – Myb - AurB network determines the choice between mitotic or endoreplication cycles (Figure 7). Each of these proteins act in a complex: CycA / CDK1 regulates mitotic entry, Myb is a subunit of the MMB transcription factor complex, and AurB kinase is one of four subunits of the CPC. While each of these complexes were previously known to have important mitotic functions, our data indicate that they are key nodes of a network whose activity level determines whether cells switch to the alternative growth program of endoreplication (Figure 7). Our results are consistent with previous evidence that lower activity of Myb promote polyploidization (DeBruhl et al., 2013; Katzen et al., 1998; Maqbool et al., 2010; Shepard et al., 2005). We also found that inhibiting cytokinesis downstream of the CPC results in binucleate cells that endoreplicate. Importantly, our genetic evidence indicates that not all types of mitotic inhibition result in a switch to endoreplication. For example, the Spc25 kinetochore protein and Polo kinase were both expressed at lower levels in devECs and iECs, but their knockdown resulted in a mitotic arrest, not a switch to endoreplication. These observations suggest that CycA / CDK1, MMB, and the CPC have principal roles in the mitotic network hierarchy and the decision to commit to mitosis or switch to endoreplication cycles.

**Figure 7.**
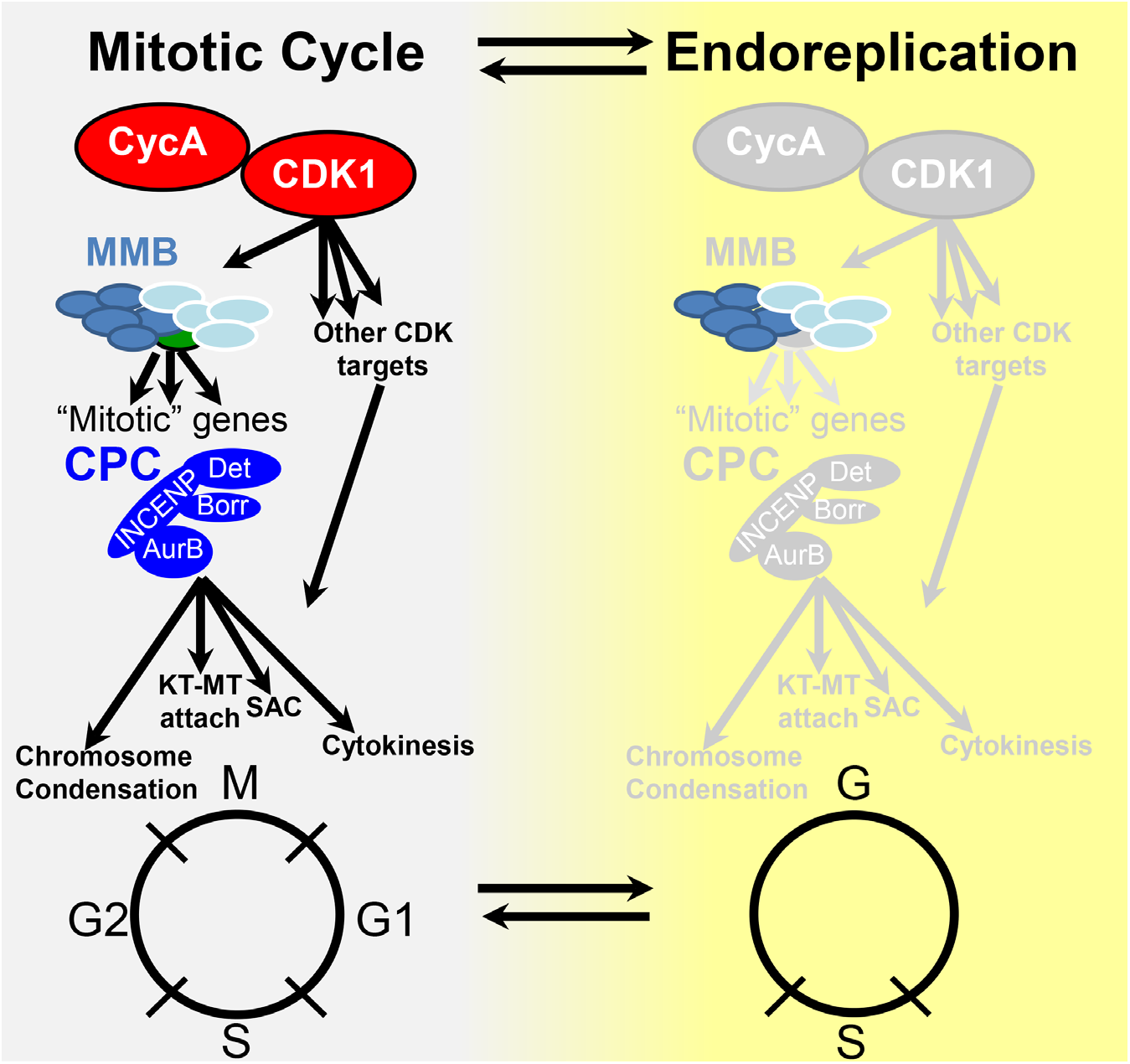
Model: The CycA – Myb – AurB network regulates the choice between cell cycle programs. Depicted are two alternative cell cycle programs, the mitotic cycle (left), and endoreplication cycle (right, yellow). During mitotic cycles, CycA /CDK1 activates the Myb-MuvB (MMB) to induce transcription of multiple genes with mitotic functions (“mitotic” genes). Among these are the subunits of the chromosome passenger complex (CPC), which phosphorylates multiple targets to regulate chromosome condensation, kinetochore-microtubule (KT-MT) attachment, the spindle assembly checkpoint (SAC), and cytokinesis. Our findings suggest that CycA / CDK1, MMB, and the CPC are key nodes of this mitotic network whose repression promotes a transition to endoreplication in both iECs and devECs. See text for further details.

While knockdown of CycA, Myb, or CPC subunits were each sufficient to induce endoreplication cycles, these iEC populations had different fractions of cells with multiple nuclei indicative of an endomitotic cycle. Knockdown of cytokinesis genes *tum* and *pav* resulted in the most endomitotic cells, followed by the CPC subunits, then *Myb*, and finally *CycA*, with knockdown of this cyclin resulting in the fewest endomitotic cells. These results suggest that knocking down genes higher in this branching mitotic network (i.e. CycA) inhibits more mitotic functions and preferentially promotes G / S endocycles that skip mitosis (Figure 7). Moreover, we found that different levels of CPC function also resulted in different subtypes of endoreplication. Strong knockdown of AurB inhibited chromosome segregation and resulted in cells with a single polyploid nucleus, whereas a mild knockdown resulted in successful chromosome segregation but failed cytokinesis, suggesting that cytokinesis requires more CPC function than chromosome segregation. It thus appears that different thresholds of mitotic function result in different types of endoreplication cycles. This idea that endomitosis and endocycles are points on an endoreplication continuum is consistent with our evidence that treatment of human cells with low concentrations of CDK1 or AurB inhibitors induces endomitosis, whereas higher concentrations induces endocycles (Chen et al., 2016). Our results raise the possibility that in tissues both conditional and developmental inputs may repress different steps of the CycA – Myb – AurB network to induce slightly different types of endoreplication cycles that partially or completely skip mitosis (Øvrebø and Edgar, 2018; Stetina et al., 2018). All together our findings show that there are different paths to polyploidy depending on both the types and degree to which different mitotic functions are repressed.

Our findings are relevant to the regulation of periodic MMB transcription factor activity during the canonical mitotic cycle. Knockdown of CycA compromised MMB transcriptional activation of mitotic gene expression, and their physical association suggests that this dependency of the MMB on CycA may be direct. While the dependency of the MMB on CycA was not previously known in *Drosophila*, it was previously reported that in human cells CycA / CDK2 phosphorylates and activates human B-Myb in late S phase, and also triggers its degradation (Charrasse et al., 2000; Saville and Watson, 1998). Unlike human cells, in *Drosophila* CycA / CDK2 is not required for S phase, and Myb is degraded later in the cell cycle during mitosis (Scaria et al., 2008; Sprenger et al., 1997). A cogent model is that CycA / CDK1 phosphorylation of Myb, and perhaps other subunits of the MMB, stimulates its activity as a transcriptional activator, explaining how pulses of mitotic gene expression are integrated with the master cell cycle control machinery (Figure 7). It remains formally possible, however, that both CycA / CDK2 and CycA / CDK1 activate the MMB in *Drosophila*. The early report that CycA / CDK2 activates Myb in human cells was before the discovery that it functions as part of the MMB, and much remains to be learned about how MMB activity is coordinated with the central cell cycle oscillator (Korenjak et al., 2004; Lewis et al., 2004).

Our data revealed both similarities and differences between iECs and devECs. Both iECs and devECs had a repressed CycA – Myb – AurB network, but, unlike devECs, iECs did not have a repressed E2F1 transcriptome. This finding was surprising because a repressed E2F transcriptome is a property shared by devECs in fly and mouse (Chen et al., 2012; Maqbool et al., 2010; Pandit et al., 2012; Zielke et al., 2011). Activator E2Fs regulate some of the same mitotic genes that are regulated by the MMB, and its downregulation may enforce a switch to endoreplication. The difference between iEC and devEC for the E2F transcriptome is relevant to the developmental control of endoreplication. The observation that both CycA^dsRNA^ iECs and devECs have lower CycA / CDK1 activity, but only devECs have lower E2F1 activity, implies that there are CDK1-independent mechanisms by which developmental signals repress the E2F transcriptome in devECs. Our findings lead us to predict that repression of the CycA – Myb – AurB network will be common among many types of iECs and devECs.

Our results have broader relevance to the growing number of biological contexts that induce endoreplication. Endoreplicating cells are induced and contribute to wound healing and regeneration in a number of tissues in fly and mouse, and, depending on cell type, can either inhibit or promote regeneration of the zebrafish heart (Cao et al., 2017; Cohen et al., 2018; González-Rosa et al., 2018; Losick et al., 2016). Determining whether the CycA – Myb –AurB network is repressed in these iECs may suggest ways to improve regenerative therapies. In the cancer cell, evidence suggests that DNA damage and mitotic stress, including that induced by cancer therapies, can switch cells into an endoreplication cycle (Davoli and de Lange, 2011; Øvrebø and Edgar, 2018; Wheatley, 2008). These therapies include AurB inhibitors, which induce human cells to polyploidize, consistent with our fly data that the CPC is a key network node whose repression promotes the switch to endoreplication (Carmena et al., 2015; Zekri et al., 2016). Upon withdrawal of AurB inhibitors, transient cancer iECs return to an error-prone mitosis that generates aneuploid cells, which have the potential to contribute to therapy resistance and more aggressive cancer progression (Chen et al., 2016). Our finding that the Myb transcriptome is repressed in iECs opens the possibility that these mitotic errors may be due in part to a failure to properly orchestrate a return of mitotic gene expression. Understanding how this and other networks are remodeled in polyploid cancer cells will empower development of new approaches to prevent cancer progression.

## Materials and Methods

### Drosophila genetics

*Drosophila* strains were obtained from the Bloomington Stock Center (BDSC, Bloomington, IN), or the Vienna Drosophila Resource Center (VDRC, Vienna Austria). The *UAS-mRFP-Myb* strain was kindly provided by Dr. Joe Lipsick. *Drosophila* were raised on BDSC standard cornmeal medium at 25°C. For the genetic screen of Figure 4, fly strains with *UAS-dsRNA* transgenes were made by the Drosophila RNAi Screening Center (DRSC) and provided by the BDSC. These strains were crossed to *dpp:Gal4*, *UAS:mRFP* and multiple progeny of each cross were scored for their adult wing phenotype. Specific details about genotypes and strain numbers can be found in supplemental data and Table S7.

### Cell Culture

S2 cells were grown at 25°C in M3 + BPYE medium supplemented with 10% Fetal Bovine Serum as described (Baum and Cherbas, 2008). iECs were supplemented with an additional 2% Fetal Bovine Serum (12% final).

### Flow Cytometry

S2 cells were treated we dsRNA for 96 hours at 25°C, harvested in PBS and fixed in ethanol. After fixation, cells were incubated in propidium iodide (20 μg/ml) supplemented with RNaseA (250 μg/ml) at 37°C for 30 minutes. Flow cytometry was performed using an LSRII (BD Biosciences) and data were analyzed with Flowjo v7.6.5 software.

### SDS-PAGE and Western Blotting

S2 cells were treated with dsRNAs for 96 hours, and total protein extracts were made. Absolute protein levels were determined by Bradford assays, and protein was separated by SDS-PAGE. Separated protein samples were electrophoretically transferred to PVDF membranes, and blotted using the following antibodies: Cyclin A (A12, DSHB), Cyclin B (F2F4, DSHB), Myb (D3R, provided by J. Lipsick), Tubulin (E7, DSHB). Blots were labeled with HRP conjugated secondary antibodies and developed using Super Signal West Pico substrate (Thermo Scientific).

### Antibody labeling and immunofluorescent microscopy

In Figure 2, S2 cells were treated with dsRNA for 96 hours at 25°C, replated on poly-D-lysine coated chamber slide, and allowed to settle for 16–18 hours. Cells were then incubated in EdU (20μM) for 2 hours at 25°C followed by click-it fluorescent labeling according to the manufacturer’s (Invitrogen) protocol. These cells were then labeled with antibodies against (pH3) (Millipore, 06–570) and appropriate fluorescent secondary antibodies. Cells were stained with DAPI (0.5μg/ml) and imaged on a Leica SP5 confocal or Leica DMRA2 fluorescent microscope. The fraction of EdU and pH3 labeled cells and nuclear area were quantified using ImageJ v1.50b software (https://imagej.nih.gov/ij/).

### iEC ovary clones

*Hsp70-FLP;Act>cd2>Gal4, UAS-mRFP* was crossed to different *UAS-dsRNA* fly strains. Well-fed 3-5 day old adult G1 females were heat induced at 37°C for 30 minutes and allowed to recover for three days before ovaries were dissected, and labeled with anti-dsRed (Takara, 632496) and anti-FasIII (7G10, DSHB) and counterstained with DAPI as previously described (Hassel et al., 2014). Cell clones in stage 6 egg chambers were imaged on a Leica SP5 confocal and Leica DMRA widefield epifluorescent microscope. Cell number was quantified by counting RFP+ cells. Nuclear area and DAPI fluorescence of individual cells within in a clone (RFP+) were measured using ImageJ and normalized to the average of wild type cells outside of the clone (RFP-) in the same egg chamber.

### RT-qPCR and RNA-Seq analysis

S2 cells were treated with dsRNA and grown at 25°C for 96 hours before RNA was harvested in TRIzol (Invitrogen) according to the manufacturer’s instructions. For RT-qPCR, cDNA was generated from 1μg RNA using the Superscript III kit (Invitrogen). qPCR was performed using Brilliant III Ultra-Fast SYBR Green qPCR Master Mix (Agilent Technologies) and the primers indicated in Table S7. Each assay was performed with technical duplicates and biological triplicates. Act5C was amplified as an internal reference control. Data were analyzed using LinRegPCR software (ver. 2016.2) the Pfaffl method to determine relative transcript levels (Pfaffl, 2001; Ramakers et al., 2003).

For RNA-Seq of S2 cells, RNA was prepared from three biological replicates of CycA dsRNA, Myb dsRNA, and GFP dsRNA treated cells. For tissues, RNA was prepared from salivary glands (SG) or brains plus imaginal discs (B-D) from the same feeding early third instar larvae in three biological replicates, as previously described (Maqbool et al., 2010). TruSeq Stranded mRNA Libraries (Illumina) were prepared by the Center for Genomics and Bioinformatics (CGB) of Indiana University according to manufacturer’s protocol. Multiplex sequencing barcodes from TruSeq RNA Single Indexes set A or B (Illumina) were added to the libraries during construction. The barcoded libraries were cleaned by double side beadcut with AMPure XP beads (Beckman Coulter), verified using Qubit3 fluorometer (ThermoFisher Scientific) and 2200 TapeStation bioanalyzer (Agilent Technologies), and then pooled. The pool was sequenced on NextSeq 500 (Illumina) with NextSeq75 High Output v2 kit (Illumina). Single-end 75 bp read sequences were generated. The read sequences were de-multiplexed using bcl2fastq (software versions 1.4.1.2, 1.4.1.2, and 2.1.0.31 for GSF1389, GSF1471, GSF1611).

### Bioinformatics

Read quality was checked with FastQC v0.11.5 (Andrews, 2010), and reads were then mapped against the Dmel R6.23 genome assembly and annotation using STAR v2.6.1a (Dobin et al., 2013). Mapped fragments were assigned to exons via the featureCounts function of the Rsubread v1.24.2 bioconductor package (Liao et al., 2013), and various psuedogenes and ncRNAs were excluded. Differential gene expression between samples was calculated using DESeq2 v1.14.1 (Love et al., 2014). Gene lists derived from RNA-Seq data sets were categorized as upregulated (Log2 fold-change ≥ 0.5 with an FDR adjusted p ≤ 0.05) or downregulated (Log2 fold-change ≤ −0.5 with an FDR adjusted p ≤ 0.05) (Benjamini and Hochberg, 1995). Human ortholog information and DIOPT scores were downloaded from FlyBase on 09-11-2018 (Hu et al., 2011) and GO terms were retrieved using the Bioconductor package AnnotationHub v2.12.0 with a snapshot date of 04-30-2018 (Morgan, 2018). GO enrichment analysis was performed and plots were generated using clusterProfiler v3.8.1 (Yu et al., 2012). The comparisons between the differentially expressed genes in the RNA-seq and the accompanying Venn diagrams were created using custom scripts and the R library VennDiagram (Chen and Boutros, 2011).

### Statistical Analysis

Statistical analysis of Figures 1B, 1C, 2B, 2G, 2H, 3B, 6K were performed using two-tailed Student’s *t* tests using Microsoft Excel (version 15.0.4753.1000). For Figure 6L, GraphPad Prism (version 7.04) was used to perform a one-way ANOVA with a two-stage linear step-up procedure of Benjamini, Krieger and Yekutieli post-hoc test (Benjamini et al., 2006) to assess statistical difference between control clones and the indicated dsRNA clones. Statistical analysis of the RNA-seq data sets from Figure 4 was done using the R packages indicated in the bioinformatics section.

## Acknowledgements

We thank D. Glover, K. McKim, J. Lipsick, and N. Perrimon for flies and antibodies. We thank E. Costello for help with the genetic screen, R Podicheti and D Rusch of the Center for Genomics and Bioinformatics (CGB) for bioinformatics support, the Drosophila Genome Resource Center (DGRC), A. Zelhof and L. Gong of the Bloomington Drosophila Stock Center (BDSC), J. Powers of the IU Light Microscopy Imaging Center (LMIC), FlyBase for critical information, and N. Perrimon and the DRSC at Harvard Medical School (NIH/NIGMS R01-GM084947) for providing transgenic RNAi fly strains. This project was supported by the Indiana Clinical and Translational Sciences Institute, funded in part by NIH #UL1TR001108, NIH R35 GM122482 to C.E.W. and by NIH R01GM113107, R01GM113107-04S1 funding to B.R.C. and C.E.W.

## Author Contributions

M.D.R. conducted all of the experiments with M.J.D. helping with RNA isolation and antibody labeling. R.A.P. performed the bioinformatics analysis. A.J.W., A.W.K. and A.M.B. conducted the knockdown screen in the wing. B.R.C and M. D. R. conceived the project and wrote the manuscript. M.A.L., C.E.W., and G.E.Z provided expert advice, experimental guidance, and helped with writing of the manuscript.

## Supplemental Data

### List of supplemental tables

**Table S1 – Differentially expressed genes in CycA dsRNA iECs**

**Table S2 – Differentially expressed genes in Myb dsRNA iECs**

**Table S3 – Shared upregulated genes in CycA dsRNA iECs and Myb dsRNA iECs**

**Table S4 – Shared downregulated genes in CycA dsRNA iECs and Myb dsRNA iECs**

**Table S5 – Shared downregulated genes in CycA dsRNA iECs, Myb dsRNA iECs and devECs (salivary glands)**

**Table S6 – Results of RNAi wing screen**

**Table S7 – Full list of fly strains and primers used**

### Supplemental Figure Legends

**Figure S1.**
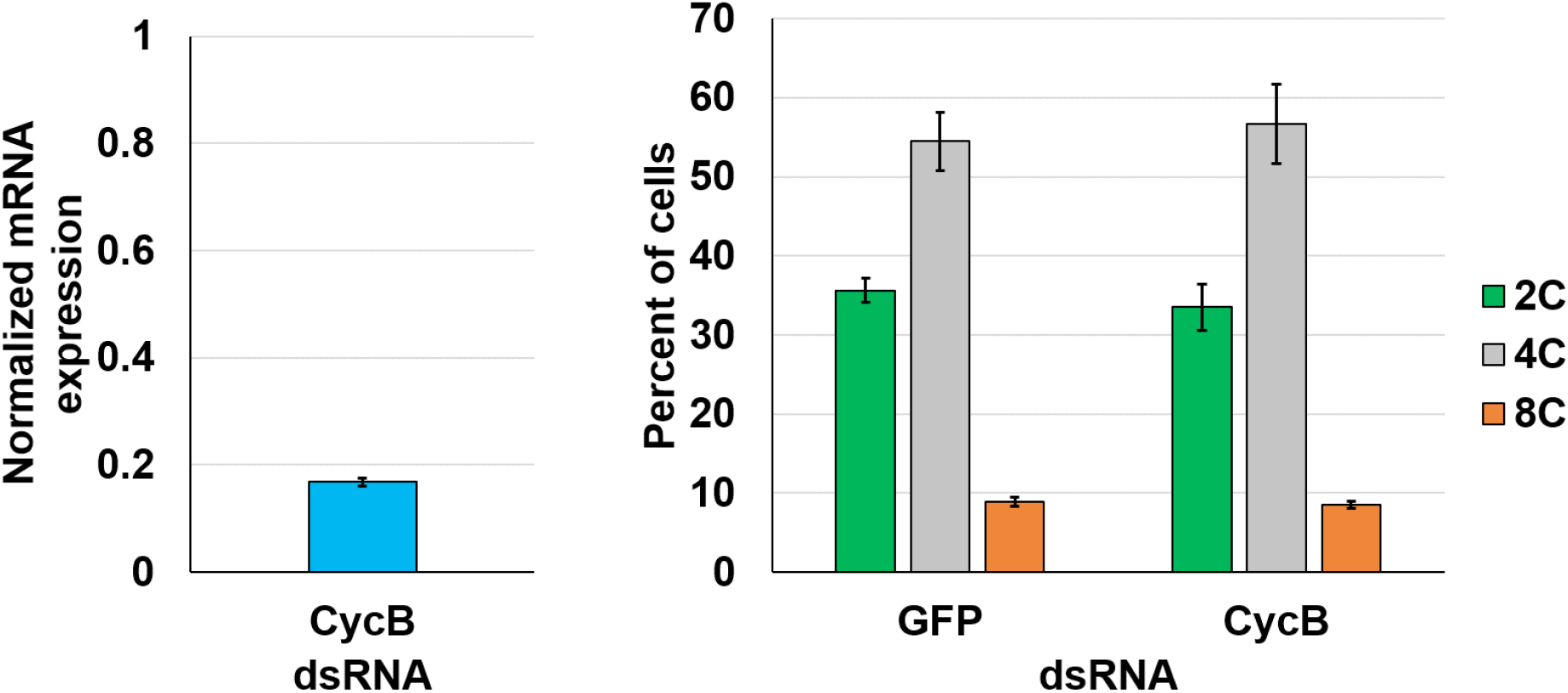
Knockdown of CycB does not induce endoreplication. S2 cells were treated with CycB dsRNA. (**A**) qRT-PCR quantification of CycB transcript in CycB dsRNA versus GFP dsRNA control cells. (**B**) Quantification of flow cytometry data for ploidy classes in GFP dsRNA and CycB dsRNA cells (mean and S.D. for N=2).

**Figure S2.**
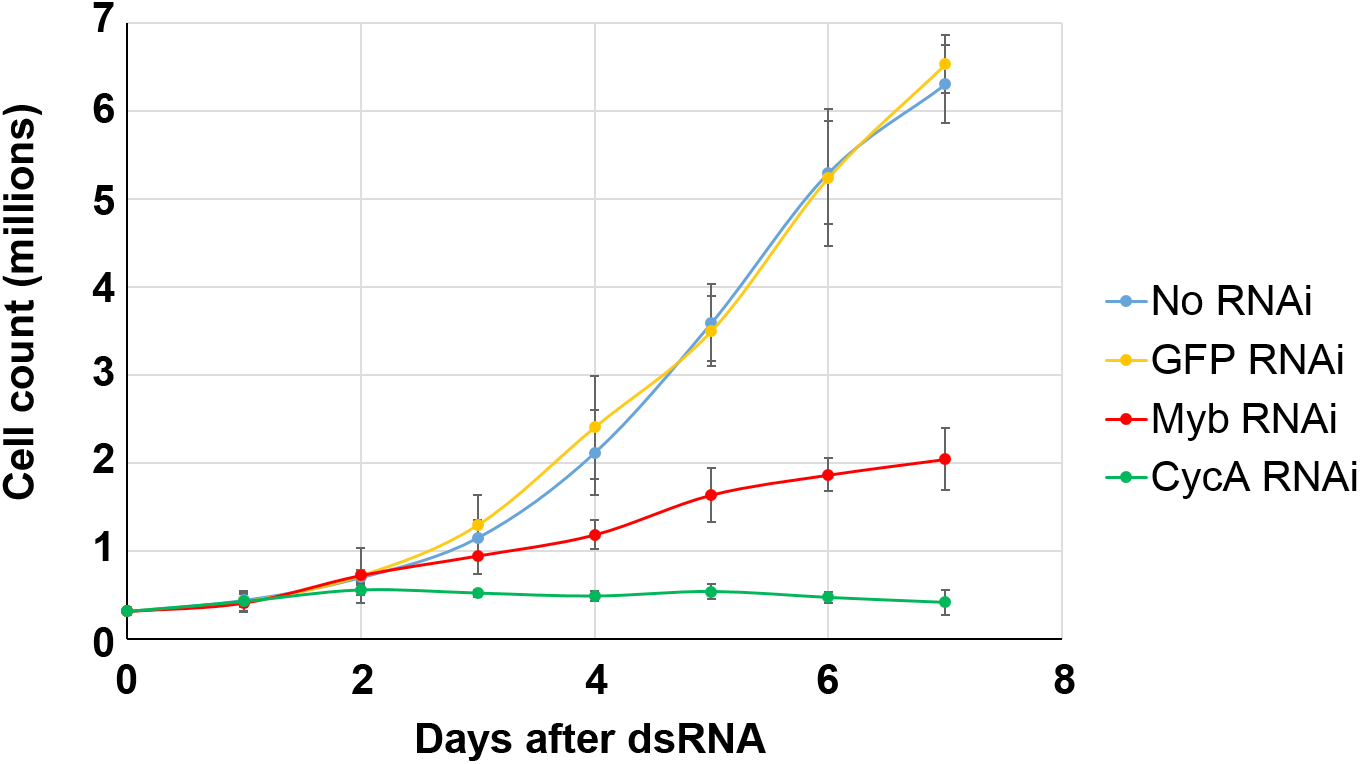
Knockdown of CycA or Myb inhibits cell proliferation. 5e05 cells were plated and treated with the indicated dsRNAs. The cells were counted once every 24h for 7 days (mean and S.D. for N=3).

**Figure S3.**
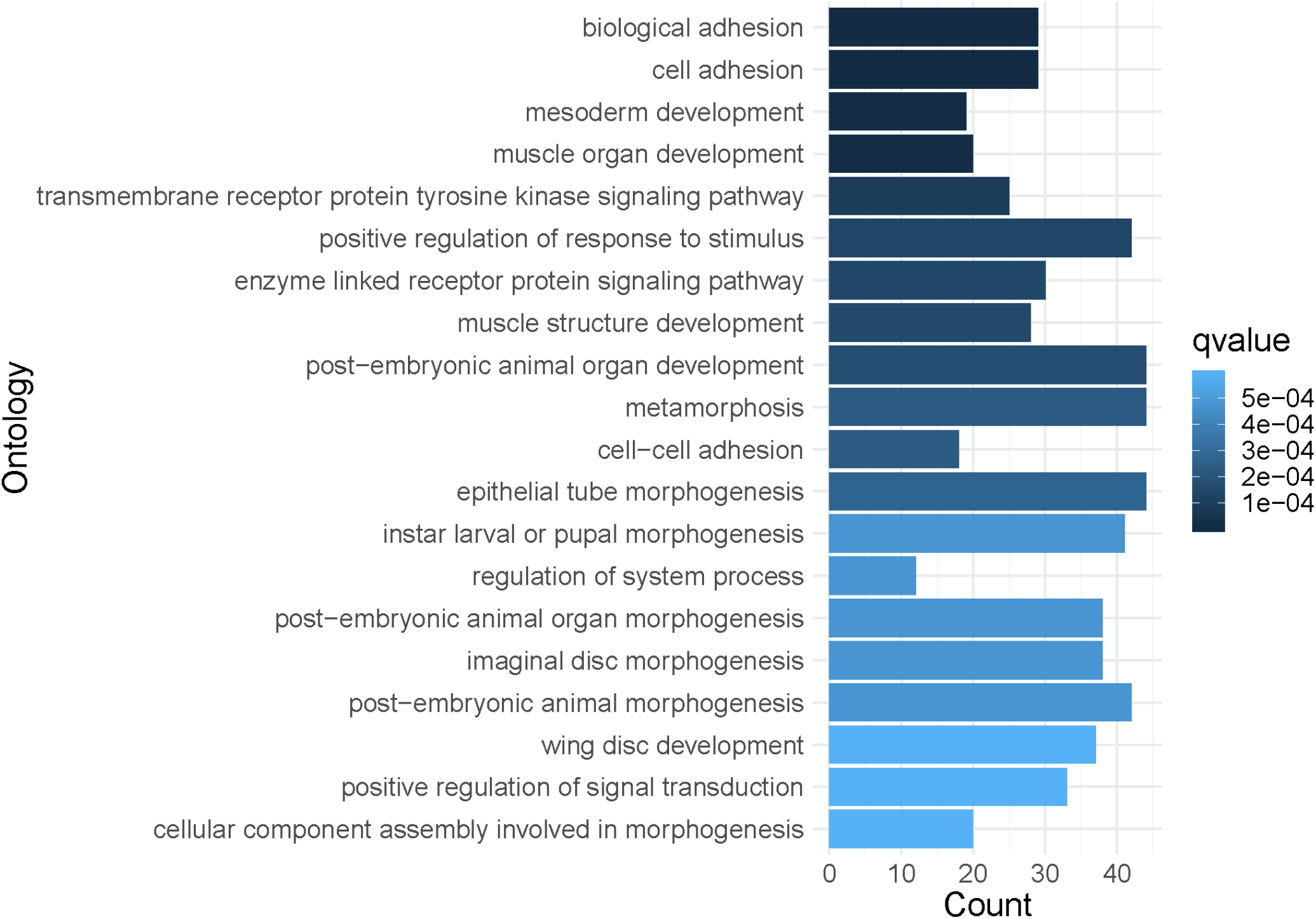
CycA dsRNA iECs and Myb dsRNA iECs have increased expression of genes involved in development and metabolism. Biological Process (BP) Gene Ontology (GO) category analysis was performed on genes that were upregulated at least Log2FC 0.5, with an FDR corrected q <0.05 in both the CycA dsRNA, and Myb dsRNA iECs relative to GFP dsRNA treated cells. The graph shows number of genes in the top 20 GO categories that were significantly enriched in both iEC types with color coding indicating FDR corrected q value for that class.

**Figure S4.**
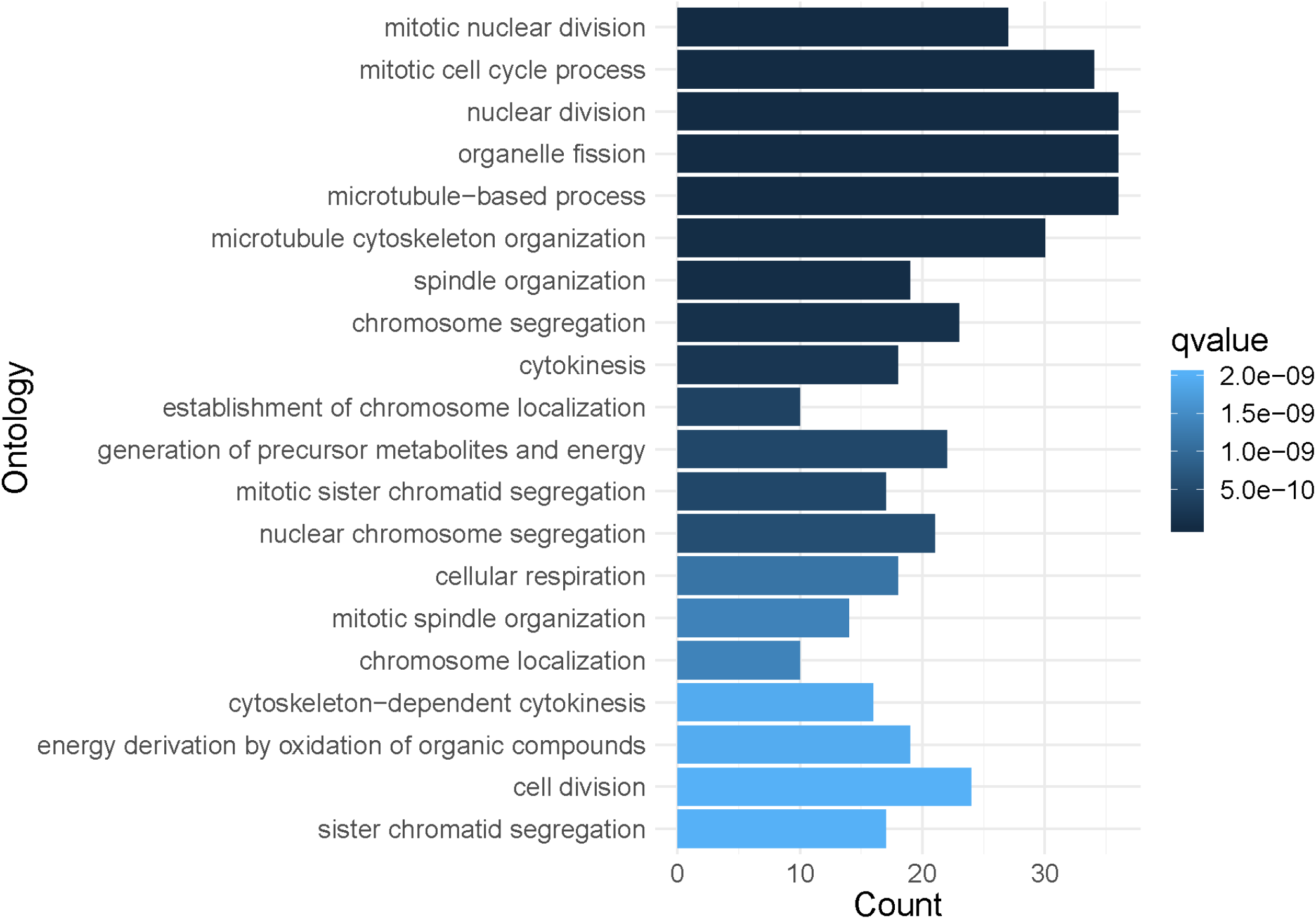
CycA dsRNA iECs and Myb dsRNA iECs have decreased expression of genes required for mitosis. BP GO category analysis was performed on genes that were downregulated at least Log2FC −0.5, with an FDR corrected q <0.05 in both the CycA dsRNA, and Myb dsRNA iECs relative to GFP dsRNA treated cells. The graph shows number of genes in the top 20 GO categories that were significantly enriched in both iEC types with color coding indicating FDR corrected q value for that class.

**Figure S5.**
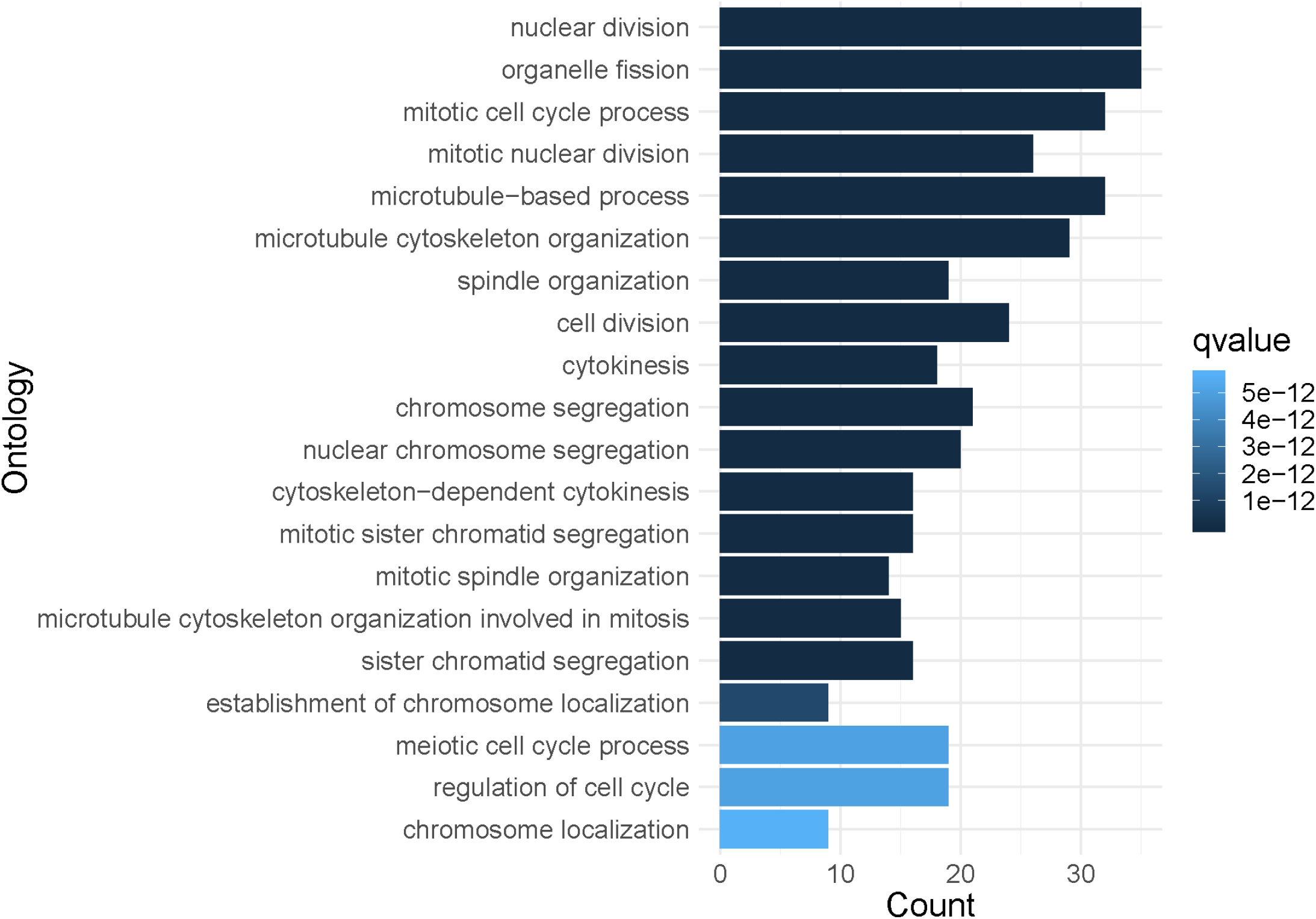
iECs and devECs have decreased expression of Myb-induced genes that are required for mitosis. Comparison of RNA-Seq results for iEC in culture and devEC in salivary glands. BP GO category analysis was performed on genes that were downregulated at least Log2FC −0.5, with an FDR of <0.05 in the CycA dsRNA, and Myb dsRNA iECs relative to GFP dsRNA treated cells and the salivary gland endocycling vs Brain-disc tissues. The graph shows the top 20 BP GO categories that were significantly enriched in the overlap of CycA dsRNA iECs, Myb dsRNA iECs, and salivary gland devECs.

